# Increased pattern similarity in two major olfactory cortices despite higher sparseness levels

**DOI:** 10.1101/2021.04.15.440031

**Authors:** Chaviva Markind, Prosenjit Kundu, Mor Barak, Rafi Haddad

## Abstract

Pattern separation is a fundamental process that enhances discrimination of similar stimuli and can be achieved by sparsening the neural activity and expanding the coding space. Odor stimuli evoke patterns of activity in the olfactory bulb (OB) and these activity patterns are projected to several cortical regions that contain larger numbers of neurons and show sparser activity levels. However, whether these projected patterns are better separated is still unclear. Here we compared odor responses in the OB, anterior piriform cortex (aPC) and anterior olfactory nucleus (AON) to the exact same odor stimuli. We found that odor representations are more similar, noisier and less distinctive in aPC and AON than in the OB. The increase in similarity was observed for both similar and dissimilar odors. Modeling odor transformation from the OB to the olfactory cortex using simulated as well as actual OB odor responses as inputs, demonstrates that the observed rise in odor representation similarity can be explained by assuming biologically plausible variation in the number of OB inputs each cortical neuron receives. We discuss the possible advantages of our findings to odor processing in the aPC and AON.

**Highlights:**

- Odor representations in the aPC and AON are more correlated despite increase in sparseness levels.
- Odor identity is best represented in the OB.
- Variance in the number of inputs from OB can explain the reduction in odor separation.

## Introduction

Pattern separation is the process by which similar neural representations become more distinct. Classical theoretical works suggest that reduction in pattern similarity can be achieved through expansion from a low dimensional space to a high dimensional space together with sparsening ^1^. Dimensionality expansion and sparse representations that improve pattern separation have been reported in many sensory systems, the cerebellum and in the hippocampus ^2,3^.

Expansion through random projection and sparsening are well documented in the olfactory system. In Drosophila, ~150 antenna lobe projection neurons (PNs) output olfactory information to ~2500 Kenyon cells in the mushroom body ^4^. Kenyon cells are much more selective to odors because they have higher response thresholds, receive broad feedback inhibition, integrate from a small number of PNs and decay quickly. As predicted by theory, odor ensemble representations in the mushroom body are decorrelated ^5^. Similarly, in the olfactory bulb (OB) of rodents, there are ~50,000 mitral and tufted (MT) cells that project mostly non-topographically to several olfactory cortical regions including the anterior olfactory nucleus (AON) and anterior piriform cortex (aPC) ^6–8^. These regions are believed to contain at least an order of magnitude more neurons than in the OB ^9^.

It is generally thought that odor representations in the aPC are sparser than in the OB ^9–16^ and mostly decorrelated, and that the aPC makes odor discrimination more robust ^16–19^, facilitating odor identity coding ^20,21^. A recent study that modeled the OB to PC circuitry as a random feed-forward network that expands and sparsens the input, showed that PC should decorrelate odor activity patterns ^14^. One empirical study found that odor mixtures that differ by more than one component are indeed less similar in aPC than in OB ^22,23^. However, a comprehensive comparison of odor representation similarity in OB and aPC is still lacking. Furthermore, how odors are represented in the AON is much less investigated and understood.

Here we compared odor representations and coding principles in the OB, AON and aPC using a set of nine odorants which include similar and dissimilar odor-pairs. We defined odor similarity by either representing each odorant in the physicochemical space using their molecular descriptors, or in the neural space using the OB activity patterns. Our results indicate that in contrast to what is expected from theory, odor representations in these cortical regions are more correlated, noisier and represent odor identity less distinctively than the OB. This increase in similarity occurred despite higher sparseness levels in the cortex and for odor-pairs of all similarity levels. Using simulation models that use experimental and simulated OB odor responses as inputs, we suggest a biologically plausible modification of the aforementioned feed-forward network ^14^ that can explain these findings.

## Results

### Odor representations in the OB, aPC and AON

To compare odor representations across different brain regions we extracellularly recorded the neural responses to nine odor stimuli in the OB, aPC and AON in anesthetized mice under the exact same experimental conditions (Figure 1A, Methods). The nine odorants used included structurally diverse odorants well separated in the physicochemical odor space ^24^ as well as similar ones (Figure 1B, Methods). We recorded the activity of 101 neurons in OB, 200 in aPC and 138 in AON. As in previous studies, the spontaneous activity was higher in the OB than in the cortex (Figure 1–figure supplement 1A). The spiking activities in individual neurons in the three regions were strongly coupled to respiration (Figure 1C-D and Figure 1–figure supplement 1B). Interestingly, compared to the OB and aPC, a relatively small percentage of AON neurons preferred to fire during inhalation (Figure 1D). The population mean in all three regions peaked shortly after the transition between inhalation and exhalation, with mean aPC and AON neurons’ odor-evoked peaks occurring slightly before the OB (Figure 1E and Figure 1–figure supplement 1C). Odor responses began in the three regions in the first sniff post odor onset and continued throughout the three respiration cycles that occurred during the 2 seconds of odor presentation (Figure 1–figure supplement 1C-D). As reported in previous studies, the number of odors each neuron in the OB responded to was distributed exponentially with most neurons not responding to any of the odors and very few responding to multiple odors (Figure 1F). On average, each odor activated 3.63 ± 0.617% (mean ± SE) and suppressed 2.09 ± 0.348% of OB neurons in the first sniff post odor onset (Figure 1G-H). Consistent with ^25^, very little inhibition was found in the AON (0.966 ± 0.241% suppressed vs. 19.163 ± 1.22% activated neurons) and neuron responses to odors were stronger and more broadly tuned. aPC neurons’ response selectivity was in between, more similar to OB (Figure 1F-H).

**Figure 1.**
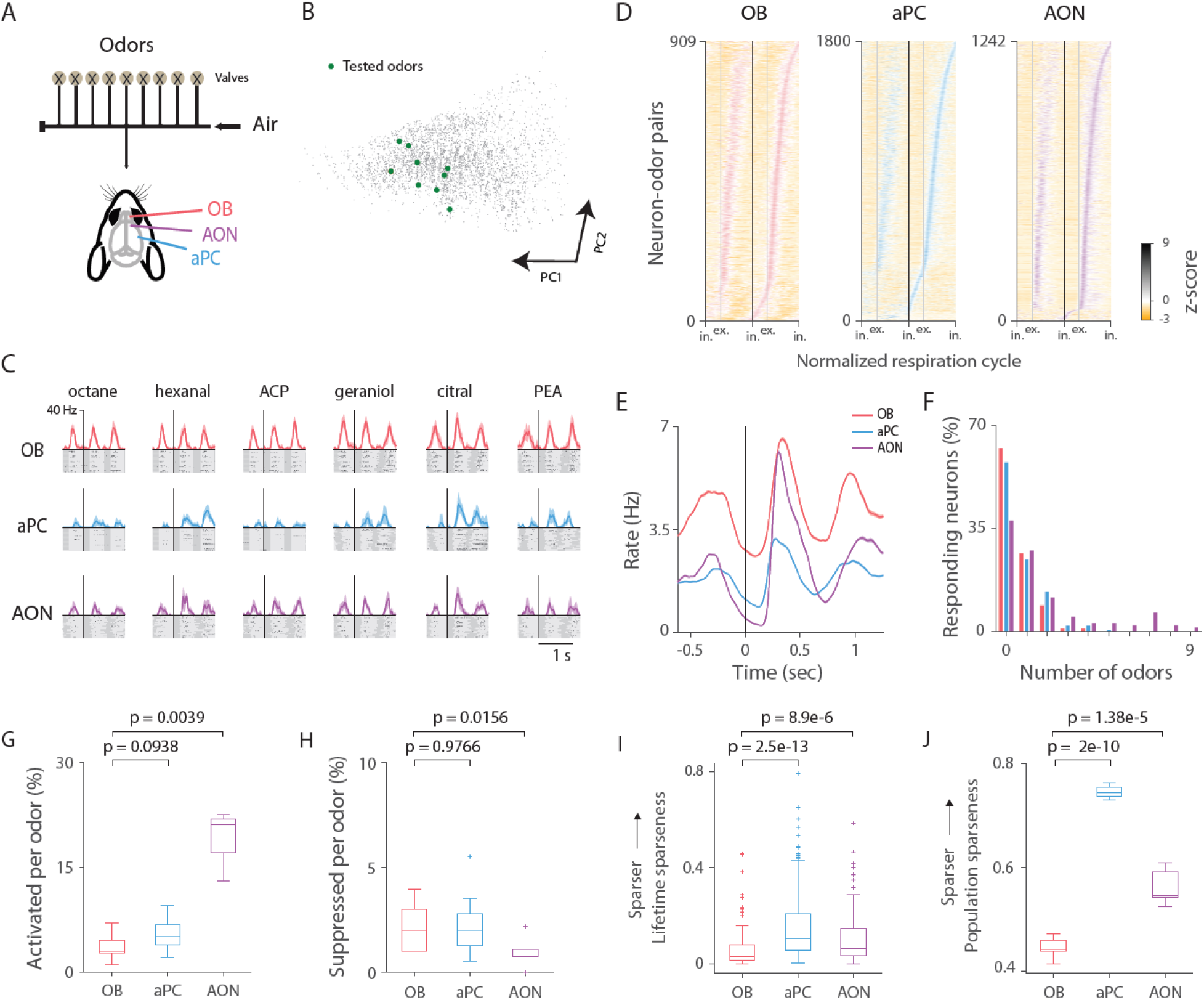
Odor-evoked activity in the olfactory bulb, anterior piriform cortex and anterior olfactory nucleus. **(A)** Experimental schematic. Extracellular recordings of neural responses to nine odor stimuli in OB, aPC and AON in anesthetized mice. **(B)** 4359 odorant molecules depicted in principal component space (Methods). Green circles mark the odorants used in the study. **(C)** Responses of example neuron in OB (red), aPC (blue) and AON (purple) to twenty presentations of six odors. See methods for all nine odorant names. Raster plots and PSTHs are aligned to the first inhalation post odor onset. Dark gray and light gray shadings indicate the inhalations and exhalations, respectively. Odor duration was 2 seconds. **(D)** Normalized PSTHs for all OB, aPC and AON neuron-odor pairs sorted by latency to peak in the first sniff post odor onset. Each respiration duration is stretched or squeezed according to the median respiration duration. Spikes were reassigned to their original relative inhalation or exhalation phase in this mutual standardized respiration cycle. ‘in.’ and ‘ex.’ indicate the inhalation and exhalation start times, respectively. Black vertical line marks the start of the first inhalation post odor onset. **(E)** Average PSTH of all odors’ mean elicited responses across all neurons (101 neurons in OB, 200 in aPC and 138 in AON, Methods). **(F)** Percentage of neurons that significantly responded to specific numbers of odors (p < 0.05, Wilcoxon rank-sum test) in the first sniff post odor onset in OB, aPC and AON. **(G-H)** Percentage of neurons per odor that responded significantly with an increase **(G)** or decrease **(H)** in spike count in the first sniff post odor onset (OB vs aPC or OB vs AON, Wilcoxon signed-rank test). **(I-J)** Treves-Rolls lifetime and population sparseness in the first sniff post odor onset (Wilcoxon rank-sum test and paired t-test (df = 8), respectively).

To compare the sparseness levels in the three brain regions, we calculated the commonly used Treves-Rolls lifetime and population sparseness indices ^26^ with modifications ^12,27^. The Treves-Rolls methods estimate the amount of non-uniformity of the neural response to the stimuli (Methods). Computing the Treves-Rolls indices we found that odor responses in both the aPC and AON are significantly sparser than in the OB in terms of lifetime and population sparseness (Figure 1I-J).

### Odor identity is best represented in the olfactory bulb

Theoretical considerations suggest that expansion of the neural space and sparsening the neural responses play a key factor in decorrelating responses in feedforward networks and improve linear separability. To analyze the similarity of different odor representations at the population level, we represented each odor as a vector of evoked mean spike counts during the first sniff post odor onset. We averaged across trials as in previous studies ^12,15,28,29^ so as to reduce the inherent trial-to-trial variability. We subtracted the baseline activity to reflect signal correlation rather than the high baseline population correlation due to spontaneous firing rates (see Methods). Using these activity vectors, we computed the Pearson correlation coefficient between all odor pairs (36 odors pairs in three brain regions, Figure 2A-B). We found that odor representations were significantly more correlated in the cortex than in OB (p = 0.00112, OB vs aPC; p = 1.63e-14, OB vs AON, paired t-test, df = 35). Odor representations in AON were particularly similar (mean ± SEM, r_AON_ = 0.6567 ± 0.025), intermediate in aPC (r_a_PC = 0.361 ± 0.025) and least similar in OB (r_OB_ = 0.234 ± 0.036). Compatible results were obtained when we used spike rates instead of spike counts (Methods). Odor representations were more similar in the cortex than in OB for odor-pairs of diverse similarity levels in the physicochemical space (Figure 2–figure supplement 1A). Notably, when comparing odor-pairs based on their similarity level in OB, in both cortical regions the increase in pairwise similarity for non-similar odor pairs was greater than for similar odor pairs (Figure 2C).

**Figure 2.**
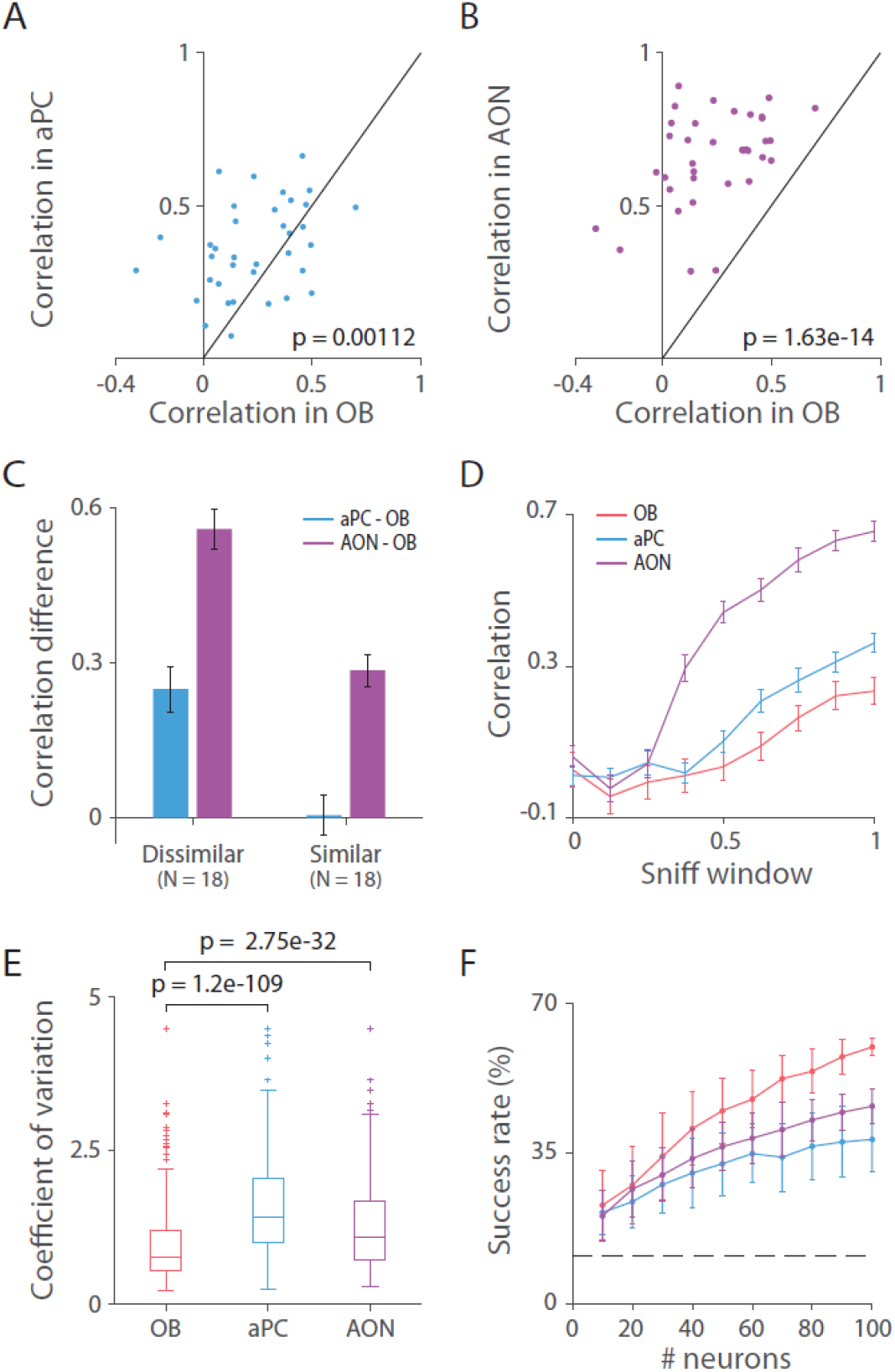
Odor identity is best represented in the olfactory bulb. **(A-B)** 36 odor-pairwise correlations in the first sniff post odor onset in aPC **(A)** and AON **(B)** versus those in OB. Identity line is indicated in black. Mean odor-evoked neural representations are more similar in the cortex than in OB (paired t-test, OB vs aPC or OB vs AON, df = 35). **(C)** Mean ± SEM of the difference in correlations in the first sniff between the cortical regions and OB of odor-pairs that are dissimilar (r < median) and similar (r ≥ median) in OB. There is a larger increase in odor similarity in the aPC and AON for dissimilar odors than for similar ones. **(D)** Mean ± SEM odor pairwise correlations calculated using the activity in accumulative windows of the first sniff post odor onset (window size is an 1/8 of the sniff). **(E)** The coefficient of variation across trials in the first sniff post odor onset. Trial variability is higher in the cortical regions than in the OB (Wilcoxon rank-sum test, OB vs aPC or OB vs AON). **(F)** Odor identification decoding accuracy as a function of the number of neurons. OB has higher decoding accuracy than aPC and AON. Displayed is the mean ± SD of the success rate of the decoder on all 9 odors in 100 random samplings of neurons (out of a total of 101, 200 and 138 neurons in OB, aPC and AON, respectively). Decoder was trained on activity vectors of the average spike counts in the duration of the three sniffs taken throughout the odor presentation, across 15 trials. Dashed line marks the chance level accuracy (11.11%).

MT neuron odor responses have complex temporal dynamics including epochs of excitation and inhibition ^30,31^ while aPC neuron odor responses are much less dynamic with typically one transient excitatory epoch ^12,19^. To test if this distinction may underlie the difference in odor similarity, we calculated the odor correlations in accumulative and moving window bins of the first sniff and found that the mean odor-pair correlations increased in aPC and AON relative to OB throughout the respiration cycle (Figure 2D and Figure 2–figure supplement 1B). The result was consistent across several sniffs post odor onset in both the aPC and the AON (Figure 2–figure supplement 1C).

The increase in odor representation correlation in aPC and AON may suggest that odor identities are better represented in the OB than in the two subsequent cortical regions we examined. However, an increase in similarity does not necessarily imply a reduction in representation separation as it also depends on the response variability. Low response variability could result in an improved representation even when distinct objects are represented as more similar to one another. We therefore compared the odor response trial variability across brain regions using the coefficient of variation metric (CV, Methods). The CV is defined as the ratio of the standard deviation to the mean and therefore is a normalized measure of variability and suited for comparison of brain regions with different baseline and mean odor response firing rates. We found that the trial variability is higher in the cortex than in OB in both aPC and AON (Figure 2E and Figure 2–figure supplement 1D).

The analyses so far show that odor representations are noisier and more similar in the two examined olfactory cortical regions. To better evaluate the level of odor representation separation in these regions, we conducted a decoding analysis. This analysis takes into account both the similarity of the representations of different odors and the variation within trials of the same odor. To this end, we tested how well a linear decoder can identify an odor given the population response of one held-out trial. We trained a centroid-based leave-one-out pattern matching linear decoder on activity vectors of the neurons’ spike counts in the sniffs during the odor stimulation (Methods). We used a linear decoder because theory suggests that expansion and sparsening are expected to improve linear separability. Performing a decoding analysis as a function of the number of neurons we found that OB has higher identification decoding accuracy than aPC and AON (Figure 2F. See Methods).

We conclude that odor representations in the OB are more decorrelated and more accurate in terms of response consistency and identity decoding than in the AON and aPC. This contrasts with the common belief that aPC neurons perform pattern separation.

### Checking our result on published data

To test whether the increase in pattern similarity and reduction in odor identity decoding that we observed is specific to our selection of odors or experimental setup, we utilized data from a recently published study of neural activity recorded simultaneously in the OB and aPC of several awake mice passively exposed to six different odors (Figure 3–figure supplement 1A, ^19,32^). Comparing OB and aPC odor representations in this dataset, we found consistent results: Odor representations in aPC are sparser than in OB (Figure 3A-B), more correlated for both similar and dissimilar odors (Figure 3C-D and Figure 3–figure supplement 1B-D) and more variable (Figure 3E). Furthermore, odor identity decoding accuracy is higher in OB than in aPC (Figure 3F). It is important to note here that our measure of odor similarity is different than the measure in ^16,18^, where correlations were computed on the trial-by-trial responses and without subtracting the baseline. We computed the correlation as in previous related studies ^12,15,28,29^ using the trial-averaged evoked responses because it is more suitable for the purpose of this study which is to compare odor representation across brain regions. Computing trial-by-trial correlations is beneficial for comparing within and between odor similarities.

**Figure 3.**
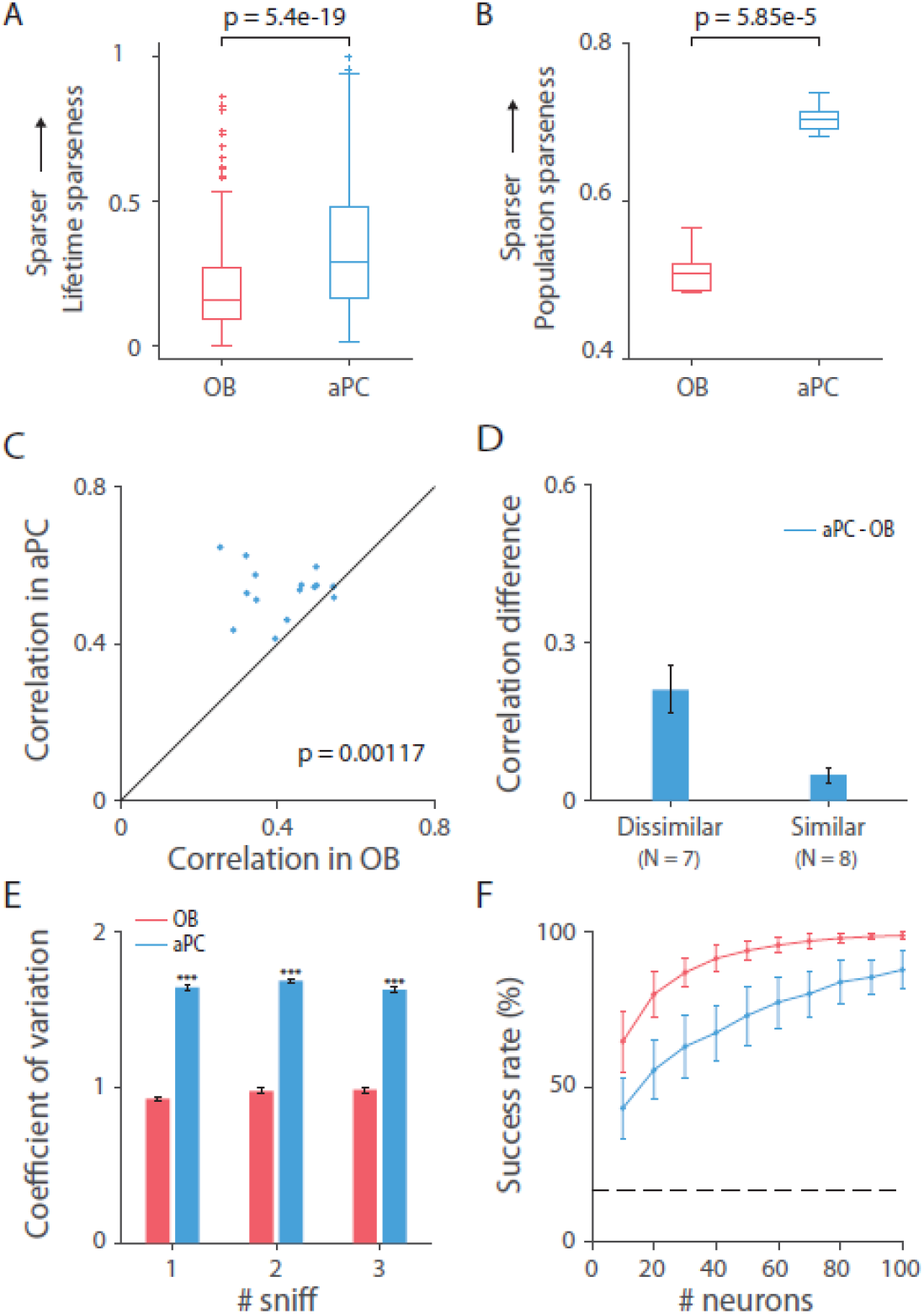
The reduction in odor representation in the cortex extends beyond our datasets. This figure displays results of analyses conducted on data of OB and aPC published in ^19,32^. **(A-B)** Treves-Rolls) lifetime and population sparseness in the first sniff post odor onset (Wilcoxon rank-sum test and paired t-test (df = 5), respectively). **(C)** 15 odor-pairwise correlations in the first sniff post odor onset in aPC versus those in OB (paired t-test, OB vs aPC, df = 14). **(D)** Mean ± SEM of the difference in correlations in the first sniff between aPC and OB of odor-pairs that are dissimilar (r < median) and similar (r ≥ median) in OB. **(E)** Mean ± SEM coefficient of variation across trials in the first three sniffs post odor onset (Wilcoxon rank-sum test for each sniff, OB vs aPC). **(F)** Odor identification decoding accuracy as a function of the number of neurons. OB has higher decoding accuracy than aPC. Displayed is the mean ± SD of the success rate of the decoder on all 6 odors in 100 random samplings of neurons (out of a total of 271 and 659 neurons in OB and aPC, respectively). Decoder was trained on activity vectors of the average spike counts in the duration of the two sniffs taken throughout the odor presentation, across 15 trials. Dashed line marks the chance level accuracy (16.67%).

### Variance in the number of inputs increases odor correlations

The experimental results show that although the odor neural response is expanded and sparsened in the aPC and AON compared to the OB, its identity representation is not improved in terms of neural representation correlation and identity decoding. This is in contrast to what is expected from a simple feed-forward transformation with random connections as was demonstrated in a recent feed-forward model that simulated a two-layer network representing the OB and PC ^14^. In this model, each of the 10000 simulated PC neurons received input from a random set of 60% of the total 1000 simulated OB neurons. Each PC neuron applied a Heaviside function on the sum of its OB inputs after subtracting a fixed threshold. The threshold was chosen such that an average of 6.2% of PC neurons were activated by the odors, matching their imaging data ^14^. It was assumed that the number of OB inhibitory inputs is twice the number of excitatory ones with half the efficacy (i.e., ~40% of the connections had a weight of −0.5 and ~20% had a weight of 1). These parameters ensured balanced excitation and inhibition inputs that mimic the balanced afferent excitatory and recurrent inhibitory inputs each aPC neuron receives ^13^. Probing this model with three sets of simulated odor responses with different levels of similarity (non-class odors, 30% overlap class odors, and 70% overlap class odors) it was shown that, odor pairwise correlations in PC are expected to be lower than in the OB (Figure 4A, dots below the identity line).

**Figure 4:**
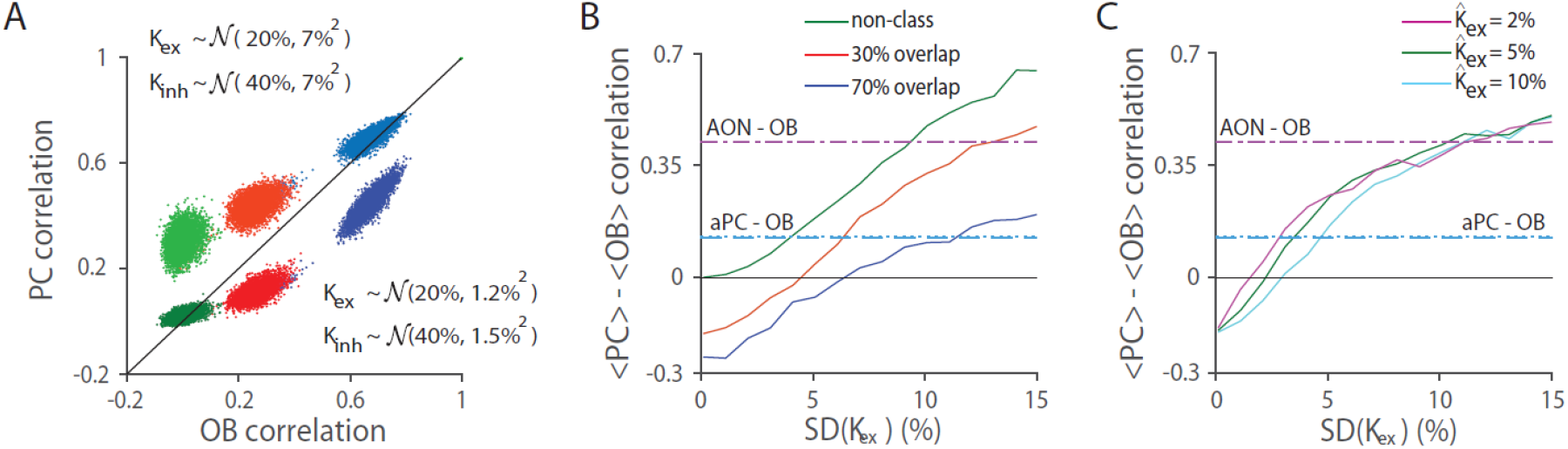
Variability in the number of connections increases odor-pairwise correlations. **(A)** Correlation of PC ensemble representations versus correlation of OB ensemble representations. Green, red and blue dots represent pairwise correlations of non-similar odor pairs (non-class), mildly similar odor pairs (30% glomeruli overlap) and highly similar odor pairs (70% glomeruli overlap) as in Schaffer et al. model. Lighter versions of the same colors are used to plot the corresponding classes when we assumed that the number of connections with positive weights (K_ex_) and negative weights (K_inh_) are drawn from normal distributions with the same means but larger variances, as specified. The values of the means and standard deviations are stated as percentages of OB neurons. **(B)** The difference between the mean correlations in PC and OB as a function of the standard deviation of the number of positive connections, with the mean number of connections fixed as in **(A)**. The variability in the negative connections is the same as in the positive connections. Blue dotted and dashed overlapping lines mark the difference between the experimental mean correlations in aPC and OB in our dataset and in the Bolding 2018 dataset, respectively. Purple dash-dot line marks the difference between the experimental mean correlations in AON and OB data. **(C)** The difference between mean pairwise correlation in PC and OB as a function of the standard deviation of the number of positive connections for the 30% overlap class of odors with different average numbers of positive connections, where the average number of negative connections is twice as large with half the efficacy. The variability in the negative connections is the same as in the positive connections. Colored horizontal lines as in **(B)**.

We sought to understand how a feed-forward network could nevertheless underlie the experimentally observed increase in correlations and decrease in identity decoding. The above model assumed that each PC neuron integrates from normally distributed numbers of OB inputs with relatively small variability (SD of 1.2% and 1.5% for the excitatory and inhibitory connections, respectively). The number of OB neurons each aPC and AON neuron receives inputs from and their variability is currently unknown. An anatomical study estimated the number of direct inputs to aPC and AON to be up to a few 10’s ^6^ while an electrophysiological study suggested that the number of direct (and probably indirect) connections to aPC might be as high as 10% of the overall number of glomeruli ^33^. We therefore examined how odor representations depend on the variability in the number of connections for various average number of connections (keeping the ratio between excitatory and inhibitory connections set to two). First, fixing the average number of inputs as in the model described above, we found that odor representation correlation increased as a function of connection number variability (Figure 4A-B). For the non-class and 30% overlap class, assuming a SD of ~4-6% or ~9-13% of the number of OB inputs can explain the increases in odor representation correlation observed in the aPC and AON experimental data, respectively (Figure 4B). Moreover, the increase in correlations in the simulated PC was larger for odors that were considered dissimilar (less correlated) in the OB (Figure 4A), consistent with the experimental results (Figure 2C and Figure 3D). Simulating the transformation with different average numbers of connections showed that the increase in odor correlations due to variability in the number of connections was stronger when the average number of connections and the SD levels were relatively small (Figure 4C, for the 30% overlap class).

### Exploring the effect of the threshold level, number of connections and their variability on odor representations in the cortex

We next explored the effect of the parameters in a similar feed-forward model that uses actual OB neural odor responses as inputs, rather than artificially generated odor response patterns. We assumed that as in the original model, each PC neuron integrates from a random set of OB neurons and responds only if the sum of the inputs it receives from the OB is higher than some threshold value *T*. That is,

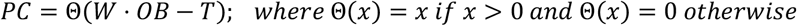

where *OB* and *PC* denote ‘neurons X odors’ response matrices of the OB and PC neurons’ odor responses, respectively; and *W* is the connectivity matrix through which PC neurons are connected to the OB neurons. We used the 271 X 6 OB neuron-odor trial-averaged responses from the Bolding 2018 dataset as this dataset was larger and therefore more suitable for this simulation, but the results are similar when using the OB neurons from our recordings as input to the model. To remove the high inherent correlation caused by the spontaneous firing of MT cells and to take into account that some odors actually reduce the number of spikes an MT neuron fires, and therefore the number of spikes the PC neuron receives, we subtracted their baseline activity (Methods). Thus, in this model MT responses could have positive and negative values reflecting excitatory and inhibitory responses, respectively. We therefore set all connection weights to one. We set the number of PC neurons to be ten times the number of available OB neurons.

We first examined how PC neurons’ threshold level affects PC odor representations as a function of the number of inputs each PC neuron receives from the OB, when there is no variability in the number of inputs. To assess this, we no longer assumed that only 6.2% of PC neurons respond on average and that excitation balances inhibition. We first noticed that when responses were only required to be non-negative (i.e., T = 0), the odor correlations in PC did not differ from those in the OB, regardless of the number of connections (pink line in Figure 5A). The sparseness level decreased as we increased the number of connections (Figure 5B, pink line). Increasing the threshold level confirmed that PC odor representations were decorrelated and sparsened (Figure 5A-B), as expected by theory ^34^ and as demonstrated in ^14^. Increasing the number of connections counteracts the effect of thresholding as the PC activity becomes less decorrelated and its sparseness level begins to saturate (Figure 5A-B). Considering threshold values that are drawn from a normal distribution revealed stronger decorrelation and sparsening effects (Figure 5C-D).

**Figure 5.**
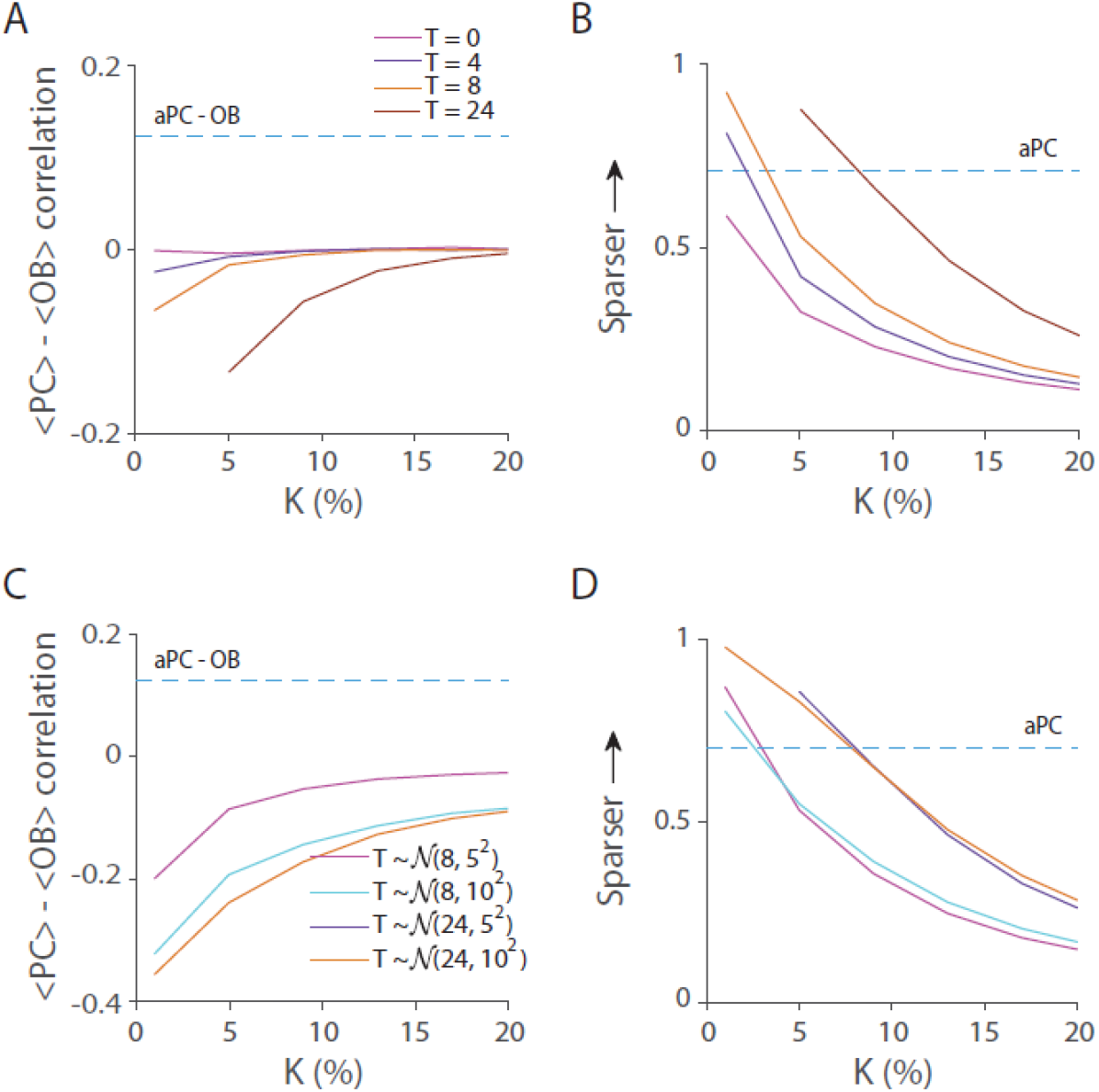
Thresholding decorrelates and sparsens odor responses. **(A)** The difference between PC and OB mean odor pairwise correlations as a function of the number of connections (marked as K) for several selected threshold values. The mean correlation did not change in the absence of a shift from zero thresholding *(T = 0*, pink line). Higher thresholding values decorrelate PC odor representations when the number of connections is relatively small. Blue dashed horizontal line marks the mean difference in the Bolding 2018 dataset. **(B)** Population sparseness level of the simulated PC neuron responses for the threshold values in **(A)**. Thresholding sparsened the PC neural activity for small K. Blue dashed horizontal line marks the mean population sparseness in the aPC of the Bolding 2018 dataset. **(C)** Mean odor pairwise correlation difference when the threshold (T) is normally distributed as specified. (**D**) Population sparseness levels for each threshold distribution corresponding to **(C)**. Blue dashed lines as in **(A-B)**.

We next examined how the variability in the number of connections affects odor representations. We found that as in the previous model, the higher the variability the more correlated the odor representations became (Figure 6A). The actual mean connection number did not have a substantial effect on the increase in odor correlations (Figure 6A) but did affect the sparseness level, with high number of connections tending to generate a denser neural response (Figure 6B). When assuming the number of connections is distributed according to an exponential distribution, we observed an increase in correlation and decrease in sparseness levels as the mean number of connections (and therefore the standard deviation) increased (Figure 6C-D).

**Figure 6.**
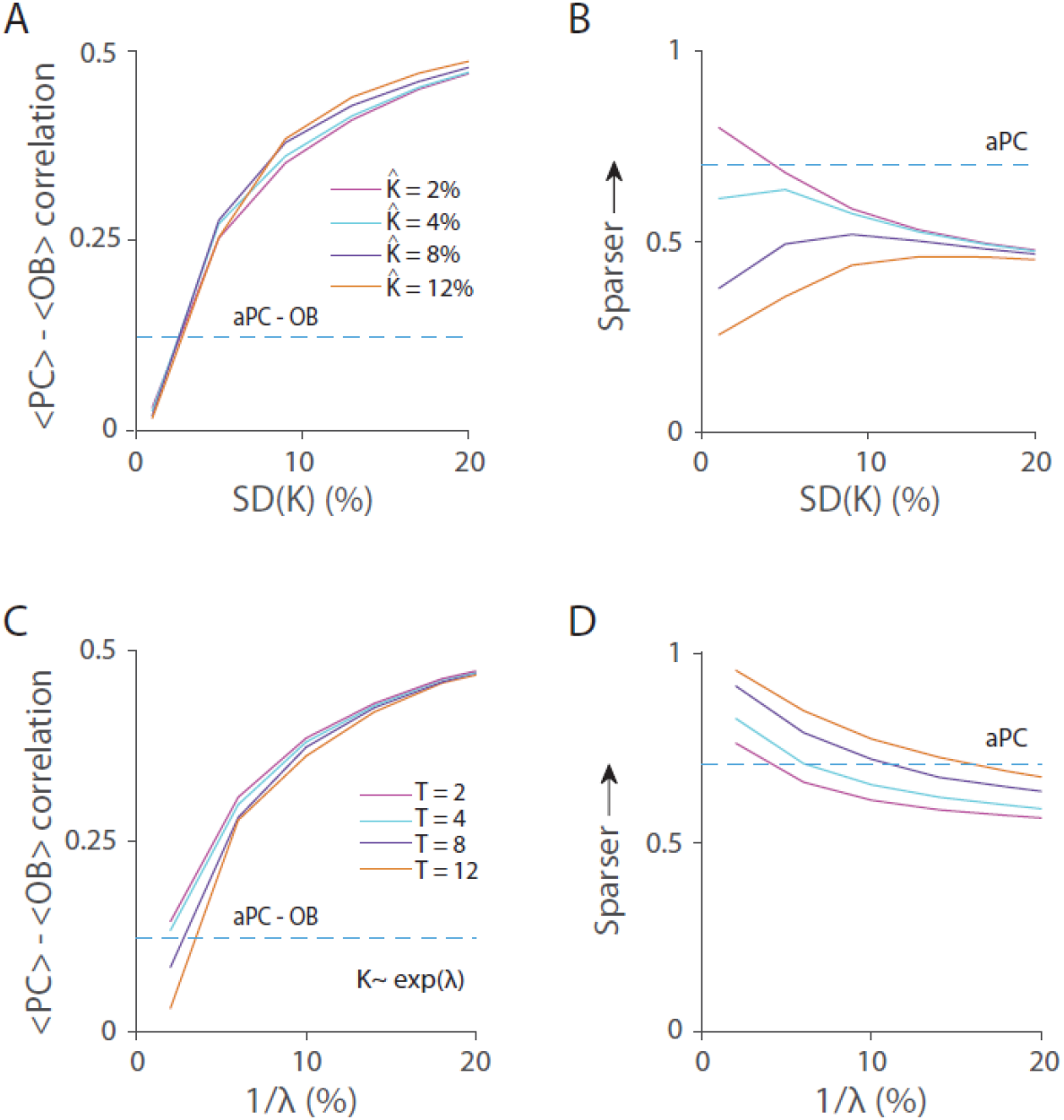
Variability in the number of PC inputs increases odor pairwise correlations in PC. **(A)** Mean difference in correlation as a function of the standard deviation of the number of connections (marked as K). Several mean K values were examined, as specified. Threshold was fixed to 12 spikes/sniff. The odor-pair correlations increase as the variability in K increases, irrespective of the mean number of connections. **(B)** Population sparseness levels corresponding to the conditions in **(A)**. Blue dashed lines as in Figure 5. **(C-D)** Same as in **(A-B)** when the number of connections is extracted from an exponential distribution as specified by λ. Results are displayed as a function of the average number of connections, which is equal to the standard deviation of the distribution. The correlation increases and sparseness decreases as the SD of the number of connections increases.

Finally, we assessed the effect of variability in the number of connections on odor identity decoding. For this purpose, we input to our model the OB responses without subtracting the baseline, as was done for the decoding analysis of the experimental data (see Methods), and randomly set inhibitory and excitatory weights as in Schaffer’s model (see Methods). We performed a decoding analysis using the simulated PC neurons as inputs to the decoding algorithm. We found that consistent with the increase in correlations, the decoding accuracy decreased with the increment in connection variability in both the normal and exponential distribution models, irrespective of the average number of connections or threshold values (Figure 7A-B).

**Figure 7.**
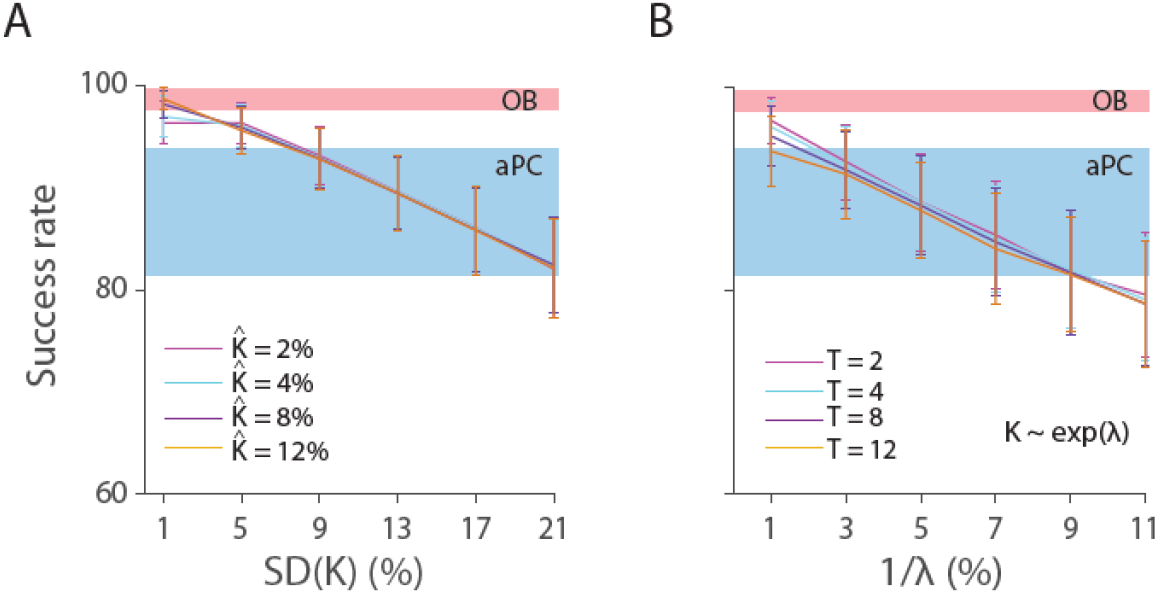
The effect of variability in the number of connections on odor decoding accuracy. **(A-B)** Odor identification decoding accuracy as a function of the variability in the number of connections. Decoding accuracy decreases with the increase in variability. Results show the mean ± SD of the success rate of the decoder on six odors of 20 different simulations of PC, where in each simulation the mean was calculated from 50 random samplings of 100 PC neurons out of a total of 2710. Red and blue patches mark the decoding accuracy range (mean ± SD) computed when using 100 neurons in OB and aPC from the Bolding 2018 dataset. **(A)** The number of connections is drawn from a normal distribution as specified. The threshold was fixed to 12 spikes/sniff. **(B)** The number of connections is distributed exponentially as specified. Several threshold values were examined.

We conclude that assuming a reasonable level of variability in the number of connections each PC neuron forms with the OB can explain the observed increase in odor representation correlations and decrease in identity decoding accuracy, despite higher sparseness levels. Specifically, assuming that each PC neuron is randomly connected to ~1-5% of the OB inputs and that these numbers follow an exponential distribution, or a normal distribution with low SD fits well with the experimentally observed increase in odor pairwise correlations, reduced decoding accuracy and increase in sparseness levels in aPC.

## Discussion

### Expansion, sparseness and separability

Increasing coding space through expansion and sparsening of the neural activity and thresholding are considered key computational mechanisms underlying the improvement in neural representation observed in several sensory systems, the cerebellum and the hippocampus ^1^. Our results show that although information from the OB is expanded and sparsened in the aPC and AON, its representation is not improved in terms of similarity and identity coding. We further verified the results using data collected in another lab using different odors and a different awareness state. These findings may reflect a fundamental distinction between odor coding and other sensory stimuli coding. A visual stimulus, for example, is decomposed into basic features such as contrast and colors, which are then integrated to build more complex features such as orientations, and these are further integrated to robustly represent the object invariantly to scale and rotation ^35^. Odors, on the other hand, are encoded by the subset of receptors that are activated by the odorant molecules. Moreover, odor responses are decorrelated by several OB interneurons that further improve odor representations ^28,29,36–38^. This may suggest that unlike in some other sensory systems, no additional decorrelation stages are required as the OB already has all the necessary odor information to robustly identify the odor.

During the process of writing this manuscript two papers have been published that examined odor representations in the OB and PC. Bolding et al. ^16^, also found a reduced decoding accuracy in aPC compared to the OB in the awake state as we found in our datasets and in our analysis of the data from ^19^. However, in the anesthetized state they found that odor identity decoding accuracy is actually better in aPC than in OB. One possible reason for this difference in result is the anesthesia methods used. We started the recording at least an hour after the first anesthesia induction and administrated additional doses of anesthesia as the animal showed signs of awakening. In Bolding et al, a single bolus of injection was given, and the experiments were conducted shortly after initial induction within the first ~30-45 minutes. This might ensue a different level of anesthesia which results in stable neural activity in aPC as they found. This might also explain why they observed a surprising reduction in OB and an increase in aPC responsiveness during anesthesia which was not found in previous OB ^39–41^ and PC studies ^15^. We note that this study also reported a reduced trial-to-trial correlation in aPC. As stated above (and see Methods) we used the trial-averaged evoked activity to estimate odor representation similarity as it is more suitable to the analysis conducted here. Trial-to-trial correlation depends on the response variability and thus the reduction in trial-to-trial correlation they found is consistent with the increase in aPC response variability and the lower odor identity decoding accuracy in the aPC that we found.

A second study by Pashkovski et al. ^15^, reported that odor correlations in PC were higher for chemically similar odors and reduced for dissimilar ones in awake artificially breathing mice. In our and in the Bolding 2018 datasets, we find that dissimilar odors as quantified by their OB representation or their molecular descriptors are more similar in aPC (and AON) (Figure 2A-C, Figure 3C-D, Figure 2–figure supplement 1A, and Figure 3–figure supplement 1B). This difference between the results could be due to different recording locations. We recorded in the anterior parts of the PC while Pashkovski et al. recorded in the anterior parts of the posterior PC. The posterior PC is known to have more associative connections compared to the aPC ^20,42,43^ and therefore is considered an associative structure whereas the AON and aPC more resemble feedforward networks. Consistent with this, it has been shown that posterior PC encodes odor quality while aPC encodes odor structure ^44–46^. Interestingly, despite these differences in the correlation structure, Pashkovski et al. also found an increase in variability and decrease in decoding accuracy in this part of the PC compared to the OB as we found in AON and anterior PC.

Taken together, all these studies strongly suggest that during the transformation from the OB to the AON and the aPC, odors are represented as more similar, have higher noise levels and are harder to decode. This change in representation is in contrast to what is expected from classical random feedforward models that expand and sparsen the representation.

### Anterior PC versus OB odor coding

One possible explanation for the decrease in odor identity representation in aPC might be that aPC neurons are involved in encoding behavioral relevance. Several studies found evidence that aPC neurons are involved in valence coding ^47^, odor preference learning ^48^, appetitive odor retrieval ^49^ and flavor conditioning ^50^. Two highly dissimilar odors can have the same behavioral relevance and therefore could be encoded similarly in aPC. This may explain why identity representation is reduced in aPC compared to the OB as in all the analyzed databases, the mice did not have to attribute valence to the presented odors and the behavioral outcome is the same (i.e., ignore). However, when there is a need to differentiate between highly similar odors, aPC neurons are able to change the odor representation such that their pattern of activities are decorrelated ^22,23,51,52^.

Another possible explanation for the reduction in odor identity representations might be that the encoding ability of aPC is compromised on account of its additional functions. Recent studies found that aPC neuron-odor responses are concentration invariant ^10,18^. This may suggest that as visual object representations are invariant to scale and rotation in higher visual brain regions, odor representations become invariant to concentration at the piriform cortex. It is possible that there is an unavoidable tradeoff between identity coding and concentration invariance in the olfactory system and that the much larger number of neurons in the aPC compensates for this reduction in coding accuracy.

### AON versus OB odor coding

Little is known about the exact roles of the AON, the first area to receive direct input from the OB. OB neurons’ projections to the AON preserve some coarse spatial organization ^6^ but the functional principles underlying how AON neurons read the OB patterns remain unknown. Haberly has suggested that OB glomeruli respond to specific molecular features while AON is tuned to a specific combination of these molecular features ^53^. This interesting hypothesis suggests that AON neurons should be more selective to odors than OB neurons. However, our result, which is also supported by a previous study ^25^, found that AON neurons are relatively widely tuned with a larger number of neurons responding to each odor (Figure 1F-G and Figure 1–figure supplement 1E).

Several studies have suggested that AON may be involved in odor localization by comparing and sharing odor information between the two hemispheres ^54–58^. Neurons that are more responsive to odors irrespective of their identities may be better suited for comparing left and right inputs because they have higher chances of responding to the left or right odors.

### Learning and variability in the number of connections

Our simulation suggests a simple and biologically plausible explanation for the observed increase in similarity and decrease in odor identity representation. We show that assuming a small number of connections with some variability that is consistent with previous anatomical and electrophysiological studies can explain the observed results. What is the possible advantage of increasing connection number variability? Several studies have emphasized plasticity in the PC, pointing to a role in associative learning and experience-dependent odor recognition ^15,59–64^. One intriguing possibility is that variability accelerates learning. This has been demonstrated in recurrent neural networks ^65^ and might point to a more general mechanism. Assuming all aPC neurons have the same threshold, aPC neurons that integrate from many OB neurons with equal weights will respond even if only a subset of their inputs is active, whereas aPC neurons with a small number of connections will respond only when most of their inputs are active. The highly connected neurons will tend to respond to many odors, and this will increase the similarity in neural representation due to large numbers of shared responding neurons. This process alone is beneficial as long as the odors have no specific association as it will result in ‘built in’ generalization. However, when an odor is associated with an outcome, Hebbian learning can strengthen the active connections and weaken the non-active ones. This procedure will reduce the number of effective connections of these neurons. At the same time, the effective connection number of neurons with sparse connections will not change substantially, since for them, receiving input from a majority of their connections is required to elicit a response. The reduction in effective connection number of the neurons that have high numbers of connections will result in reduced variability in the number of effective connections across all neurons. Reducing the variability would result in decorrelating the odor responses (Figs. 4,6) and facilitate discrimination. Since high variability increases similarity, learning can be accelerated because it only needs to change the representation of one stimulus out of many similar ones clustered in one region of the neural space.

The exact number, distribution and weights of OB inputs each aPC neuron integrates from and how these values change during learning is currently unknown. Future studies that will reveal these values will shed important insights on how odors are represented and how learning shapes neural networks.

## MATERIALS AND METHODS

### Mice

All surgical and experimental procedures were conducted in accordance with the National Institutes of Health Guide for the Care and Use of Laboratory Animals and the Bar Ilan University guidelines for the use and care of laboratory animals in research and were approved and supervised by the Institutional Animal Care and Use Committee (IACUC). 15 wild-type male and female mice aged 3-6 months were used. The animals were housed in a group cage and received no experimental treatment. Animals were maintained in a reverse light/dark cycle and all experiments were performed during their dark cycle.

### Surgical Procedures

Mice were first anesthetized with ketamine/medetomidine (60/0.5 mg/kg, I.P.) and then fixed in a stereotaxic frame. The bone overlying the dorsal OB, the AON or anterior PC was removed. Additional anesthesia was administered as needed (30% of the original dose of ketamine/medetomidine). The animal’s body temperature was maintained at 36-37**°**C using a homeothermic blanket system (Harvard Apparatus).

### In vivo electrophysiology

The spiking activity of neurons was recorded extracellularly using custom built four or eight tetrodes. Neural signals were recorded using 32 channel recording system (Digital Lynx SX, Neuralynx) filtered at 300–6,000 Hz, sampled and recorded at 32 kHz and stored on a computer. Spike signals were sorted offline using spike3D. Clusters with >2% of ISIs violating the refractory period (<2 ms) were manually removed from the dataset. Neurons were recorded from the dorsal and ventral olfactory bulb. To record from the AON, we inserted the electrodes 1.25 mm laterally, −2.6 mm from bregma and 2.2 mm ventrally. To record from the anterior PC, we inserted the electrode −2.1 mm from bregma, 1.7 mm laterally and ~3 mm ventrally. A total of 101 OB neurons were recorded in 5 mice, 200 aPC neurons in 7 mice and 138 AON neurons in 3 mice. We recorded respiration using a piezoelectric sensor (APS4812B-LW100-R, PUI Audio). For verification of our results we used the data published at ^32^ of simultaneous recordings of OB and aPC. Reported are six odors and ‘blank’ mineral oil presented to awake head fixed mice. We used the sessions in which each of the six odors and ‘blank’ were presented 15 times. This dataset contains the responses of 271 neurons in OB and 659 in aPC from 10 simultaneous recordings.

### Odorants

Odorants were applied using a custom built olfactometer. Odorants were diluted in mineral oil (1:100) and stored in sealed glass vials. Odorants were delivered through a manifold converging all odor tubes into one tube that was placed in front of the animal nostrils at a distance of 1 cm. Clean air constantly flowed through the converging port to reduce cross-contamination. Airflow was controlled with a mass flow controller (Agilent, Alimc-2LSPM) and set to 0.8 l/m. Odor stimulation times and sequences were controlled by a custom MATLAB script. Odor delivery time occurred at any phase of the respiration cycle, as in natural settings. Odor stimulation duration was two seconds, with an inter-trial-interval of 20 seconds of clean air. The odor sequence was randomized, and each odor was delivered 20 times. Odorants were purchased from Sigma Aldrich at the highest possible purity. The nine odorants used were:

**Table 1.**
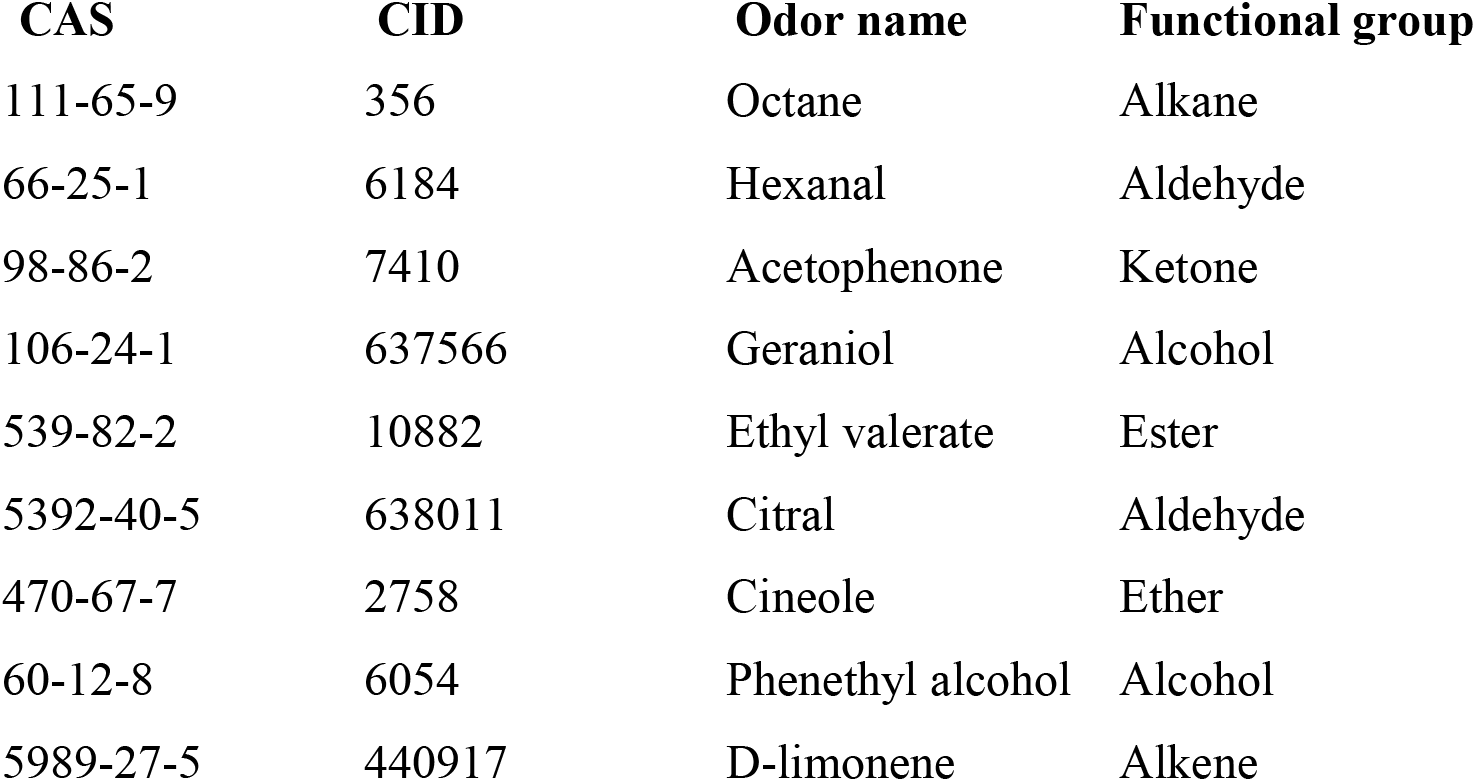

| CAS | CID | Odor name | Functional group |
| --- | --- | --- | --- |
| 111-65-9 | 356 | Octane | Alkane |
| 66-25-1 | 6184 | Hexanal | Aldehyde |
| 98-86-2 | 7410 | Acetophenone | Ketone |
| 106-24-1 | 637566 | Geraniol | Alcohol |
| 539-82-2 | 10882 | Ethyl valerate | Ester |
| 5392-40-5 | 638011 | Citral | Aldehyde |
| 470-67-7 | 2758 | Cineole | Ether |
| 60-12-8 | 6054 | Phenethyl alcohol | Alcohol |
| 5989-27-5 | 440917 | D-limonene | Alkene |

#### Analysis

### Odorants in physicochemical space

Odorants were represented using 1664 molecular descriptors (Dragon 5, Talete) and plotted in PCA space after scaling across odors as in ^24,66^. The physicochemical similarity between any two odors was computed using the Pearson correlation between the odor representations in the 1664 physicochemical space.

### Post-stimulus time histograms

To visualize trial-averaged firing rates aligned to the odor start time, spike times were convolved with a Gaussian filter with a SD of 20ms. We defined the odor start time as the first inhalation post odor onset. Most of the odor stimulations were initiated during the exhalation phase. However, when an odor stimulus was initiated during the last 50ms of the inhalation, we defined the odor start time as the following inhalation event. To estimate the population mean odor-evoked response, we averaged the PSTHs of the individual neurons for each odor, and then further averaged the nine resultant mean PSTHs. To allow for a fair comparison across sniffs, we also displayed the PSTHs with standardized inhalation and respiration cycle durations equal to the median inhalation and respiration durations in the dataset (200.8 and 628.3ms, respectively) and reassigned spike times according to their original relative inhalation or exhalation phase. To visualize the phase-locking distributions, we used this standardized respiration cycle to compute PSTHs normalized by z-score (for PSTHs that were not all-zero) for each neuron-odor pair and then sorted the neuron-odor pairs according to their latency to peak in the first respiration cycle post odor onset.

### Neural responses

Evoked responses were calculated as the mean spike count in a sniff window post odor onset (specified in each analysis), subtracted by the mean spike count in the equivalent sniff window prior to the odor onset. For the analysis of the data from ^19,32^ we calculated the odor evoked responses using the equivalent sniff window post ‘blank’ onset as the baseline activity since awake mice modulate their sniffing frequency when expecting an odor and during odor sampling.

We used spike count and not spike rate because it does not require to divide the number of spikes by the respiration duration which increases response variability due to inaccuracies in estimating the exact respiration duration. That said, the results of this study remained the same when we used spike rates.

Significant responses to odors were defined if the odor-elicited spike counts were significantly different than the respective baseline activity, according to the Wilcoxon rank-sum test (p < 0.05).

Odor neural responses depend on many factors such as, odorant identity, its concentration, flow rate and volume, vacuum position and strength, and distance from the mouse nostril. Therefore, comparison of correlation and sparseness values obtained in one study could be completely irrelevant for another study. Only comparisons between datasets collected under the exact same conditions are meaningful.

### Sparseness

The response sparseness was measured according to the Treves-Rolls sparseness index ^26^. The Treves-Rolls methods estimate the amount of non-uniformity of a neural response to the stimuli. A neuron that does not respond at all or that responds similarly to all stimuli is regarded as uniform, while a neuron that responds to only a small number of stimuli is considered sparse or selective. We modified the measure so that a value of one will indicate maximal sparseness and zero maximal uniformity ^27^. The sparseness calculation was further scaled such that it will range between 0 to 1, independent of the number of samples ^12^. The formula of the sparseness index is thus defined as follows: 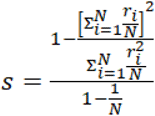. Lifetime sparseness was computed for each neuron, where N is the total number of odors and *r_i_* is the neuron’s response to odor *i*. Population sparseness was computed for each odor, where N is the total number of neurons and *r_i_* is the response of neuron *i* to the odor. Sparseness was calculated using the spike counts in the first sniff post odor onset. We used the spike count without subtracting the baseline as suggested in ^67^.

### Odor neural representation pairwise similarities

Pairwise similarities between odor representations in each brain region were evaluated by computing the Pearson correlation coefficient between activity vectors of the neurons’ average evoked spike counts across all trials. Averaging over trials reduces noise so that the computed correlation more reliably reflects the signal correlation ^12,15,28,29^. We used the evoked activity because otherwise the correlation may reflect the strong similarity in baseline activity that is very common in neurons in all olfactory regions. For example, two odors that elicited a response in a small number of non-overlapping neurons may still have a very high correlation value due to the strong resemblance of spontaneous activities in the non-responding neurons. To compare the increase in similarity in the cortex for dissimilar odors and similar odors, the odor pairs were regarded as dissimilar or similar according to their Pearson correlation in the OB or in the physicochemical space.

To calculate the Pearson correlations in windows of the first sniff, we considered the baseline activity in the anesthetized database to be the mean spike count in the equivalent window of the last sniff prior to odor onset. To calculate the evoked activity in the window bin ending in time ‘0’ (i.e., the last 1/8 of the last sniff prior to odor onset), we subtracted the mean spike count in the equivalent bin in the second to last sniff prior to odor onset. For the awake database, the baseline activity that was subtracted from the mean spike count in some sniff window of the odor trials, was the mean spike count in the equivalent sniff window of the blank trials.

### Trial variability

Odor-elicited response trial variability was assessed for the spike counts in the first few sniffs post odor onset. The variability was quantified using the coefficient of variation (CV), which is defined as the ratio of the standard deviation to the mean across trials. The results displayed in Figure 2,3 and Figure 2–figure supplement 1 are based on all neuron-odor pairs apart from those with mean spike count of zero. Results were qualitatively consistent (aPC > AON > OB) when excluding neuron-odor pairs with mean spike count lower than 1. The coefficient of variation was chosen over the Fano-factor since it is a dimensionless measurement and therefore suitable for comparing the response variability in regions with different means.

### Decoding analysis

Odor identity decoding accuracy was estimated using a centroid-based leave-one-out linear decoder. The decoder was trained on activity vectors of neurons’ spike counts in the duration of the number of sniffs taken throughout the odor presentation (3 sniffs for our dataset, 2 for the Bolding 2018 dataset). We used the neurons’ spike counts as a measure for neural activity because it reflects the number of spikes a downstream region would receive (similar results were obtained when we used the evoked spike counts). Centroid vectors were calculated for each odor as the neurons’ mean responses across trials. Since a small percentage of the neuron-odor pairs in our recordings had a few invalid trials, we used the first 15 valid trials out of the 20 to be consistent across neuron-odor pairs. For the Bolding 2018 dataset all 15 trials were used. The mean response for the test odor was computed by excluding one trial. The decoder then classified the left-out trial as the odor with the closest centroid, according to the Euclidean distance. The decoding was performed using varying numbers of neurons, where the decoding accuracy for each number of tested neurons was estimated by the mean ± SD of 100 repetitions of randomly selected neurons. In each repetition, the accuracy was calculated as the percent correct in 100 classifications of each odor.

### Modeling

In the first part of our modeling analysis we examined how changing the parameters of the previously established Schaffer et al. model ^14^ affects odor representation. The Schaffer model is a feedforward network that is based on a linear transformation and rectification using a threshold *T:*

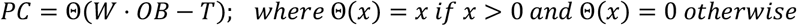

where *OB* and *PC* represent matrices of ‘neurons X odors’ responses of the OB and PC neurons, respectively and *W* is the connectivity matrix of size (# neurons in PC X # neurons in OB). Following this transformation, the PC responses were normalized to the 99th percentile value. The threshold was chosen such that an average of 6.2% of PC neurons were activated by each odor, matching their imaging data ^14^. The number of OB and PC neurons was set to 1000 and 10000, respectively. The rate of odor-activated OB neurons was set to 10% of the number of OB neurons. The OB response magnitudes of the responding neurons were taken from a multivariate lognormal distribution. The Schaffer et al. model examined three simulated odor classes: Non-class odors are a group of odors that activated non-overlapping sets of glomeruli. 30% and 70% overlap classes share 30% and 70% of the active glomeruli, respectively.

Each PC neuron was assumed to integrate from a random number of excitatory OB inputs that were normally distributed with a mean of 200 (i.e., 20%) and a SD of 12 (i.e., 1.2%). Each PC neuron was also assumed to receive counter-balancing inhibitory inputs. This was modeled by assuming each PC neuron integrates from a number of OB neurons that were normally distributed with a mean of 400 (i.e., 40%) and a SD of 15 (i.e., 1.5%) with negative weights. The weights W_ij_ of the inhibitory inputs were set to −0.5 and the weights of the excitatory connections were set to 1.

To examine how the number of connections or their variability may affect odor representation, we varied the mean number of connections and the values of the SD of the normal distributions from which the numbers of inputs were drawn. We kept the ratio between the number of inhibitory and excitatory connections to two.

In the second part of our analysis, we modified and extended the Schaffer et al. model in several ways. Instead of using simulated OB inputs from a specific predetermined distribution, we used actual trial-averaged OB data as inputs. We used the Bolding 2018 dataset because it has a larger number of OB neurons, however, the results were similar when we used the data from our OB dataset. The number of aPC neurons that were simulated was ten times the number of available OB neurons, to reflect expansion. We first considered the odor-evoked OB data in the first sniff post odor onset as inputs to the model. Since we subtracted the baseline activity, our OB responses had both negative and positive values reflecting inhibitory and excitatory responses, and we therefore assumed PC neurons integrate from all OB neurons with equal weights, W_ij_ = 1. To examine how threshold values affect odor representation we removed the assumption that on average only 6.2% of the PC neurons responded to odors and tested several threshold values. The odor-evoked OB data allowed us to examine the effect of the transformation on odor correlation and the corresponding sparseness levels. We varied the number of connections between the PC and the OB assuming either a rectified normal or an exponential distribution. The number of connections was always a positive integer.

To examine the decoding accuracy of the modeled data, we simulated PC neurons by transforming the OB data according to the proposed model and calculated the decoding accuracy of the simulated neurons. For this analysis, we used the original OB spike counts in the first two sniffs post odor onset without subtracting the baseline, consistent with the decoding analysis of the experimental data; however, congruent results were obtained when we used the OB evoked spike count. As in Schaffer’s model, we set the number of inhibitory connections to be twice as many as the excitatory ones, with half the efficacy (weights of −0.5 vs 1). We considered the OB data (neurons X odors X trials), then transformed each trial with the same connectivity matrix, whose number of connections are chosen from a rectified normal or an exponential distribution, to calculate the PC responses data (neurons X odors X trials). With this PC data we estimated the mean decoding success rate of 100 neurons by averaging over the success rate of 50 sets of 100 randomly chosen neurons from the total 2710. This procedure was repeated 20 times with different choices of connectivity matrices drawn from the same probability distributions, and we display the mean and SD of the success rate of these 20 iterations (Figure 7).

### Statistical tests

For comparing two normal distributions, the Student’s t-test (two-sided) for independent samples or paired samples was used. For comparing two independent distributions when normality cannot be assumed, significance was assessed by using the two-sided Wilcoxon rank sum and signed-rank tests. Standard error of the mean (SEM) was reported when we estimated the standard deviation (SD) of the sample mean. SD was reported when the mean was estimated from a bootstrap process.

### General experimental design

Blinding and sample size estimation are not relevant in this study and therefore were not conducted. Randomization was performed in all related experiments and analyses.

## Data and Code availability

All data and code are posted to Dryad (https://doi.org/10.5061/dryad.h18931zkf) and Github (https://github.com/rafihaddad/)

## Acknowledgments

We thank K. Bolding and K. Franks for providing the data of the awake mice and for fruitful discussions. We thank O. Barak and S.R. Datta for commenting on the manuscript. This study was supported by a grant from the I-CORE Program of the Planning and Budgeting Committee and The Israel Science Foundation [51/11 and 204/17].

## Author contribution

MB collected the data. CM, MB and RH conceived the idea and performed the analysis. PK conducted the model simulations. CM, PK and RH wrote the paper.

## Supplementary figures

**Figure 1–figure supplement 1.**
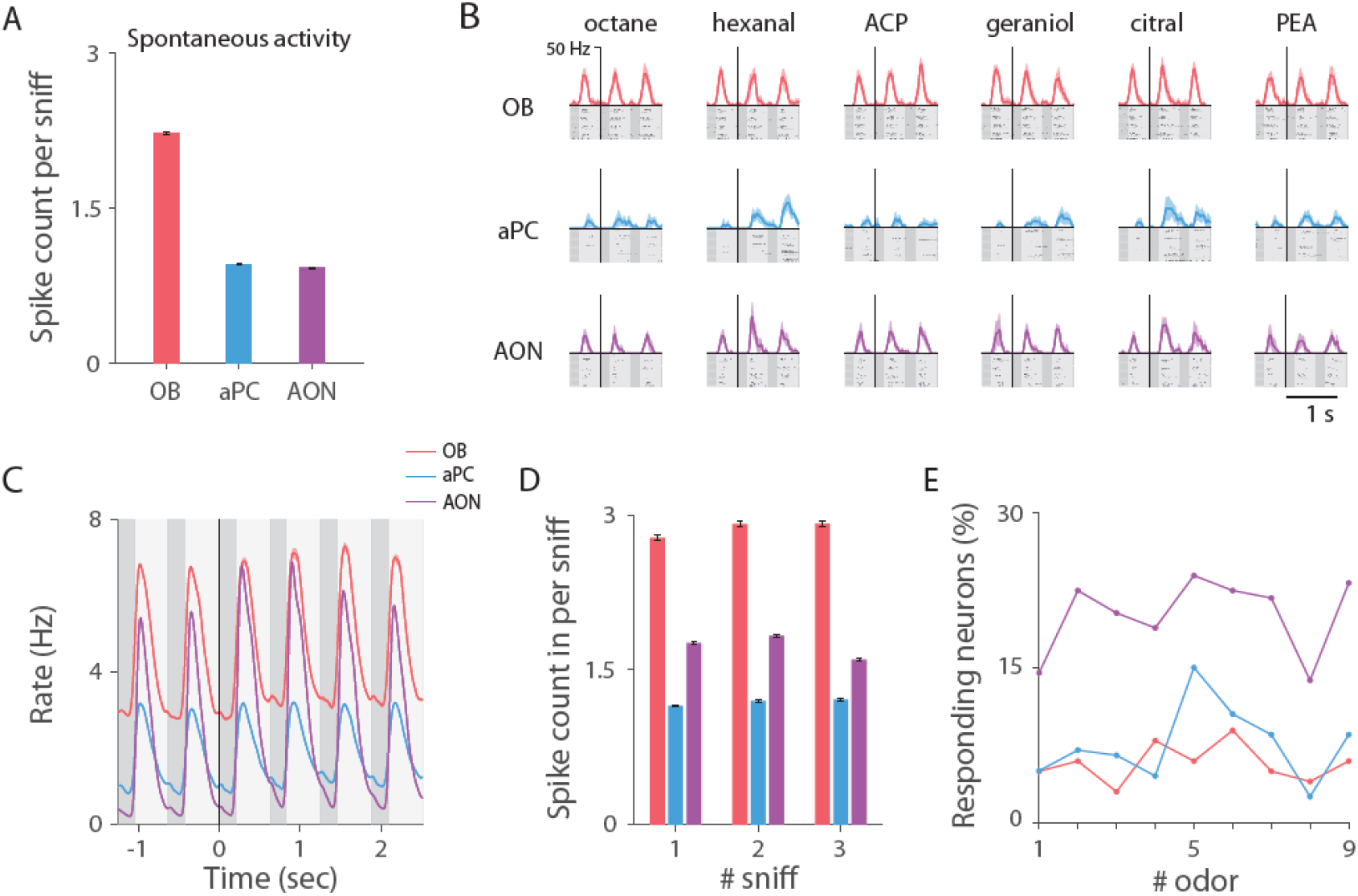
Odor-evoked activity in the olfactory bulb, anterior piriform cortex and anterior olfactory nucleus. **(A)** Mean ± SEM of the spontaneous spike count per sniff (assessed in the seven sniffs prior to odor onset across all trials). **(B)** Same examples as in Fig. 1C when the respiration cycle is standardized, as described in Fig. 1D. Raster plots and PSTHs are aligned to the first inhalation post odor onset. Color code as in Fig. 1C. **(C)** Average PSTH of all odors’ mean elicited responses across all neurons, when respiration cycle is standardized as in Fig. 1D. **(D)** Mean ± SEM of the spike count in the first three sniffs post odor onset, across all trials of all odors. **(E)** Percentage of neurons that significantly responded to each of the nine odors in the first sniff post odor onset (p < 0.05, Wilcoxon rank-sum test). AON neurons are more odor responsive than OB and aPC neurons.

**Figure 2–figure supplement 1.**
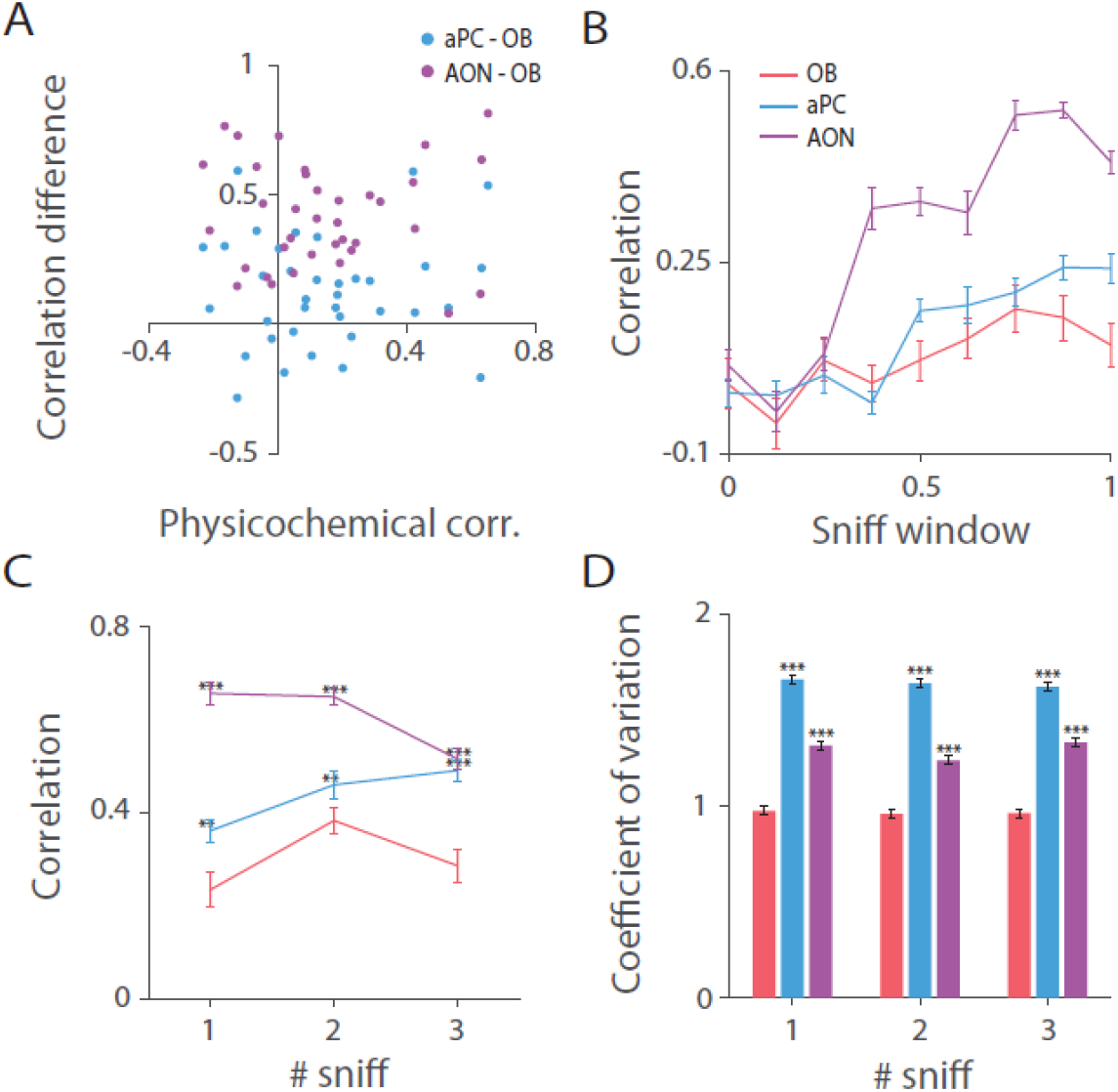
Odor identity is best represented in the olfactory bulb. **(A)**. 36 odor-pair correlation differences in the first sniff between the cortical regions and OB versus correlations computed in the physicochemical space (Methods). Odor similarity increases in the cortex compared to the OB for odor pairs of all similarity levels. **(B)** Mean ± SEM odor pairwise correlations computed in moving windows of the first sniff post odor onset (window size is an 1/8 of the sniff). **(C)** Mean ± SEM of odor pairwise correlations across the first three sniffs post odor onset. Odor pairwise similarity levels are consistently higher in the aPC and AON than in the OB. Significant differences are indicated above the cortical regions, marked by: * p < 0.05, ** p < 0.01, *** p < 0.001 (paired t-test for each sniff, OB vs aPC or OB vs AON, df = 35). **(D)** Mean ± SEM coefficient of variation across trials in the first three sniffs. Significant differences (Wilcoxon rank-sum test for each sniff, OB vs aPC or OB vs AON) are marked as in **(C)**.

**Figure 3–figure supplement 1.**
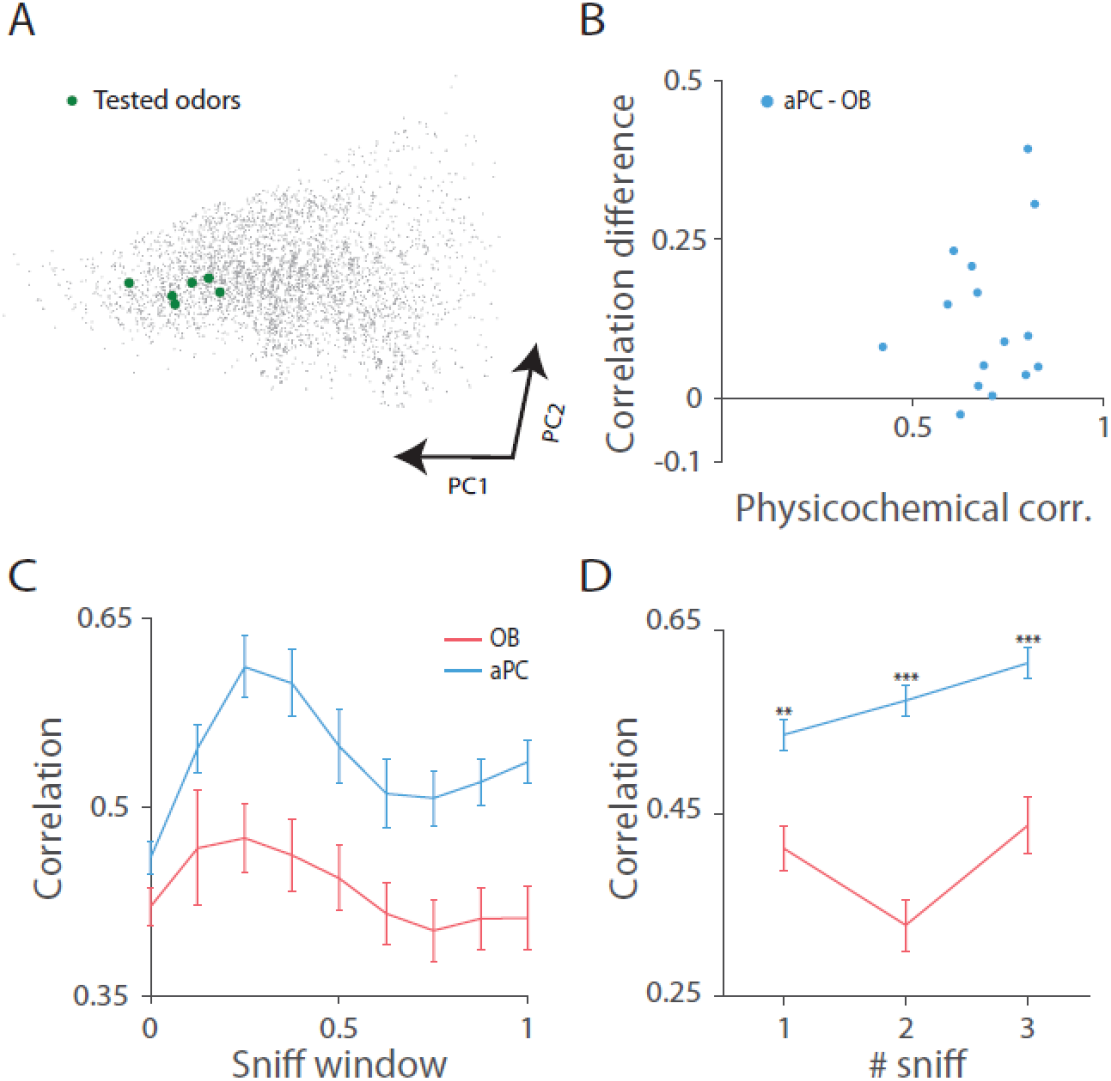
Data of OB and aPC published in (Bolding and Franks, 2018; Bolding Kevin A. Franks Kevin M., 2018). **(A)** 4359 odorant molecules depicted in principal component space (Methods). Green circles mark the odorants used in the study. **(B)** 15 odor-pair correlation differences in the first sniff between the aPC and OB versus correlations computed in the physicochemical space (Methods). Odor similarity increases in the cortex compared to the OB for odor pairs of differing similarity levels. **(C)** Mean ± SEM odor pairwise correlations calculated using the activity in accumulative windows of the first sniff post odor onset (window size is an 1/8 of the sniff). **(D)** Mean ± SEM of odor pairwise correlations across the first three sniffs post odor onset (paired t-test for each sniff, OB vs aPC, df = 14).

## Notes

### Competing Interest Statement

The authors have declared no competing interest.

## References

1. Cayco-Gajic, N. A. & Silver, R. A. Review Re-evaluating Circuit Mechanisms Underlying Pattern Separation. (2019). doi:10.1016/j.neuron.2019.01.044

2. Yassa, M. A. & Stark, C. E. L. Pattern separation in the hippocampus. Trends Neurosci. 34, 515–525 (2011).

3. Chacron, M. J., Longtin, A. & Maler, L. Efficient computation via sparse coding in electrosensory neural networks. Curr. Opin. Neurobiol. 21, 752–760 (2011).

4. Keene, A. C. & Waddell, S. Drosophila olfactory memory: single genes to complex neural circuits. Nat. Rev. Neurosci. 8, 341–54 (2007).

5. Lin, A. C., Bygrave, A. M., de Calignon, A., Lee, T. & Miesenböck, G. Sparse, decorrelated odor coding in the mushroom body enhances learned odor discrimination. Nat. Neurosci. 17, 559–568 (2014).

6. Miyamichi, K. et al. Cortical representations of olfactory input by trans-synaptic tracing. Nature 472, 191–196 (2011).

7. Sosulski, D. L., Bloom, M. L., Cutforth, T., Axel, R. & Datta, S. R. Distinct representations of olfactory information in different cortical centres. Nature 472, 213–216 (2011).

8. Kay, R. B., Meyer, E. A., Illig, K. R. & Brunjes, P. C. Spatial distribution of neural activity in the anterior olfactory nucleus evoked by odor and electrical stimulation. J. Comp. Neurol. 519, 277–289 (2011).

9. Babadi, B. & Sompolinsky, H. Sparseness and expansion in sensory representations. Neuron 83, 1213–26 (2014).

10. Roland, B., Deneux, T., Franks, K. M., Bathellier, B. & Fleischmann, A. Odor identity coding by distributed ensembles of neurons in the mouse olfactory cortex. Elife 6, 1–26 (2017).

11. Stettler, D. D. & Axel, R. Representations of odor in the piriform cortex. Neuron 63, 854–864 (2009).

12. Miura, K., Mainen, Z. F. & Uchida, N. Odor representations in olfactory cortex: distributed rate coding and decorrelated population activity. Neuron 74, 1087–1098 (2012).

13. Poo, C. & Isaacson, J. S. Odor representations in olfactory cortex: ‘sparse’ coding, global inhibition, and oscillations. Neuron 62, 850–61 (2009).

14. Schaffer, E. S. et al. Odor Perception on the Two Sides of the Brain: Consistency Despite Randomness. Neuron 98, 736–742.e3 (2018).

15. Pashkovski, S. L. et al. Structure and flexibility in cortical representations of odour space. Nature 1–6 (2020). doi:10.1038/s41586-020-2451-1

16. Bolding, K. A., Nagappan, S., Han, B.-X., Wang, F. & Franks, K. M. Recurrent circuitry is required to stabilize piriform cortex odor representations across brain states. Elife 9, (2020).

17. Srinivasan, S. & Stevens, C. F. The distributed circuit within the piriform cortex makes odor discrimination robust. J. Comp. Neurol. 526, 2725–2743 (2018).

18. Bolding, K. A. & Franks, K. M. Complementary codes for odor identity and intensity in olfactory cortex. Elife 6, 1–26 (2017).

19. Bolding & Franks. Recurrent cortical circuits implement concentration-invariant odor coding. Science (80-.). 361, (2018).

20. Bekkers, J. M. & Suzuki, N. Neurons and circuits for odor processing in the piriform cortex. Trends Neurosci. 36, 429–438 (2013).

21. Gottfried, J. A. Central mechanisms of odour object perception. Nat. Rev. Neurosci. 11, 628–641 (2010).

22. Barnes, D. C., Hofacer, R. D., Zaman, A. R., Rennaker, R. L. & Wilson, D. A. Olfactory perceptual stability and discrimination. Nat Neurosci 11, 1378–1380 (2008).

23. Wilson, D. A. Pattern separation and completion in olfaction. Ann N Y Acad Sci 1170, 306–312 (2009).

24. Haddad, R. et al. A metric for odorant comparison. Nat Methods 5, 425–429 (2008).

25. Lei, H., Mooney, R. & Katz, L. C. Synaptic integration of olfactory information in mouse anterior olfactory nucleus. J. Neurosci. 26, 12023–32 (2006).

26. Treves, A. & Rolls, E. T. What determines the capacity of autoassociative memories in the brain? Netw. Comput. Neural Syst. 2, 371–397 (1991).

27. Willmore, B. & Tolhurst, D. J. Characterizing the sparseness of neural codes. Netw. Comput. Neural Syst. 12, 255–270 (2001).

28. Gschwend, O. et al. Neuronal pattern separation in the olfactory bulb improves odor discrimination learning. Nat. Neurosci. 18, pages1474–1482 (2015).

29. Otazu, G. H., Chae, H., Davis, M. B. & Albeanu, D. F. Cortical Feedback Decorrelates Olfactory Bulb Output in Awake Mice. Neuron 86, 1461–1477 (2015).

30. Cury, K. M. & Uchida, N. Robust odor coding via inhalation-coupled transient activity in the mammalian olfactory bulb. Neuron 68, 570–585 (2010).

31. Margrie, T. W. & Schaefer, A. T. Theta oscillation coupled spike latencies yield computational vigour in a mammalian sensory system. J. Physiol. 546, 363–74 (2003).

32. Bolding Kevin A. Franks Kevin M. Simultaneous extracellular recordings from mice olfactory bulb (OB) and piriform cortex (PCx) and respiration data in response to odor stimuli and optogenetic stimulation of OB. CRCNS.org http://dx.doi.org/10.6080/K00C4SZB (2018).

33. Davison, I. G. & Ehlers, M. D. Neural circuit mechanisms for pattern detection and feature combination in olfactory cortex. Neuron 70, 82–94 (2011).

34. Wiechert, M. T., Judkewitz, B., Riecke, H. & Friedrich, R. W. Mechanisms of pattern decorrelation by recurrent neuronal circuits. Nat. Neurosci. 13, 1003–1010 (2010).

35. DiCarlo, J. J., Zoccolan, D. & Rust, N. C. How does the brain solve visual object recognition? Neuron 73, 415–434 (2012).

36. Li, W. L. et al. Adult-born neurons facilitate olfactory bulb pattern separation during task engagement. Elife 7, 1–26 (2018).

37. Yamada, Rodriguez, I. & Carleton, A. Context- and Output Layer-Dependent Long-Term Ensemble Plasticity in a Sensory Circuit. 1198–1212 (2017). doi:10.1016/j.neuron.2017.02.006

38. Chu, M. W., Li, W. L. & Komiyama, T. Balancing the Robustness and Efficiency of Odor Representations during Learning. Neuron 92, (2016).

39. Rinberg, D., Koulakov, A. & Gelperin, A. Sparse odor coding in awake behaving mice. J. Neurosci. 26, 8857 (2006).

40. Kato, H. K., Chu, M. W., Isaacson, J. S. & Komiyama, T. Dynamic sensory representations in the olfactory bulb: modulation by wakefulness and experience. Neuron 76, 962–975 (2012).

41. Kollo, M., Schmaltz, A., Abdelhamid, M., Fukunaga, I. & Schaefer, A. T. ‘Silent’ mitral cells dominate odor responses in the olfactory bulb of awake mice. Nat. Neurosci. 17, 1313–1315 (2014).

42. Hagiwara, A., Pal, S. K., Sato, T. F., Wienisch, M. & Murthy, V. N. Optophysiological analysis of associational circuits in the olfactory cortex. Front. Neural Circuits 6:18 (2012). doi:10.3389/fncir.2012.00018

43. Haberly, L. B. & Price, J. L. Association and commissural fiber systems of the olfactory cortex of the rat. J. Comp. Neurol. 178, 711–40 (1978).

44. Gottfried, J. A., Winston, J. S. & Dolan, R. J. Dissociable codes of odor quality and odorant structure in human piriform cortex. Neuron 49, 467–479 (2006).

45. Howard, J. D., Plailly, J., Grueschow, M., Haynes, J. D. & Gottfried, J. A. Odor quality coding and categorization in human posterior piriform cortex. Nat Neurosci 12, 932–938 (2009).

46. Kadohisa, M. & Wilson, D. A. Separate encoding of identity and similarity of complex familiar odors in piriform cortex. Proc. Natl. Acad. Sci. U. S. A. 103, 15206–15211 (2006).

47. Gire, D. H., Whitesell, J. D., Doucette, W. & Restrepo, D. Information for decision-making and stimulus identification is multiplexed in sensory cortex. Nat. Neurosci. 16, 991–3 (2013).

48. Morrison, G. L., Fontaine, C. J., Harley, C. W. & Yuan, Q. A role for the anterior piriform cortex in early odor preference learning: Evidence for multiple olfactory learning structures in the rat pup. J. Neurophysiol. 110, 141–152 (2013).

49. Terral, G. et al. CB1 Receptors in the Anterior Piriform Cortex Control Odor Preference Memory. Curr. Biol. 29, 2455–2464.e5 (2019).

50. Mediavilla, C., Martin-Signes, M. & Risco, S. Role of anterior piriform cortex in the acquisition of conditioned flavour preference. Sci. Rep. 6, 33365 (2016).

51. Chapuis, J. & Wilson, D. A. Bidirectional plasticity of cortical pattern recognition and behavioral sensory acuity. Nat. Neurosci. 15, 155–161 (2012).

52. Shakhawat, A. M. D., Harley, C. W. & Yuan, Q. Arc visualization of odor objects reveals experiencedependent ensemble sharpening, separation, and merging in anterior piriform cortex in adult rat. J. Neurosci. 34, 10206–10210 (2014).

53. Haberly, L. B. Parallel-distributed processing in olfactory cortex: new insights from morphological and physiological analysis of neuronal circuitry. Chem Senses 26, 551–576 (2001).

54. Esquivelzeta Rabell, J., Mutlu, K., Noutel, J., Martin del Olmo, P. & Haesler, S. Spontaneous Rapid Odor Source Localization Behavior Requires Interhemispheric Communication. Curr. Biol. 27, 1542–1548.e4 (2017).

55. Yan, Z. et al. Precise circuitry links bilaterally symmetric olfactory maps. Neuron 58, 613–624 (2008).

56. Kikuta, S. et al. From the Cover: Neurons in the anterior olfactory nucleus pars externa detect right or left localization of odor sources. Proc Natl Acad Sci U S A 107, 12363–12368 (2010).

57. Grobman, M. et al. A Mirror-Symmetric Excitatory Link Coordinates Odor Maps across Olfactory Bulbs and Enables Odor Perceptual Unity. Neuron 99, 800–813.e6 (2018).

58. Dalal, T., Gupta, N. & Haddad, R. Bilateral and unilateral odor processing and odor perception. Commun. Biol. 3, 150 (2020).

59. Choi, G. B. et al. Driving opposing behaviors with ensembles of piriform neurons. Cell 146, 1004–1015 (2011).

60. Franks, K. M. et al. Recurrent circuitry dynamically shapes the activation of piriform cortex. Neuron 72, 49–56 (2011).

61. Wilson, D. a & Sullivan, R. M. Cortical processing of odor objects. Neuron 72, 506–19 (2011).

62. Fletcher, M. L. & Wilson, D. A. Experience modifies olfactory acuity: acetylcholine-dependent learning decreases behavioral generalization between similar odorants. J. Neurosci. 22, RC201 (2002).

63. Linster, C., Henry, L., Kadohisa, M. & Wilson, D. A. Synaptic adaptation and odor-background segmentation. Neurobiol Learn Mem 87, 352–360 (2007).

64. Linster, C., Menon, A. V, Singh, C. Y. & Wilson, D. A. Odor-specific habituation arises from interaction of afferent synaptic adaptation and intrinsic synaptic potentiation in olfactory cortex. Learn Mem 16, 452–459 (2009).

65. Schuessler, F., Mastrogiuseppe, F., Dubreuil, A., Ostojic, S. & Barak, O. The interplay between randomness and structure during learning in RNNs. arXiv (2020).

66. Haddad, R. et al. Global features of neural activity in the olfactory system form a parallel code that predicts olfactory behavior and perception. J Neurosci 30, 9017–9026 (2010).

67. Rolls, E. T. & Tovee, M. J. Sparseness of the neuronal representation of stimuli in the primate temporal visual cortex. J. Neurophysiol. 73, 713–726 (1995).

